# Extraintestinal pathogenic *Escherichia coli* (ExPEC) are associated with prolonged carriage of extended-spectrum β-lactamase-producing *E. coli* acquired during travel

**DOI:** 10.1101/2020.09.23.309856

**Authors:** Boas C.L. van der Putten, Jarne M. van Hattem, John Penders, COMBAT Consortium, Daniel R. Mende, Constance Schultsz

## Abstract

**Objectives:** Extended-spectrum β-lactamase-producing *Escherichia coli* (ESBL-Ec) are frequently acquired during international travel, contributing to the global spread of antimicrobial resistance. Human-adapted ESBL-Ec are predicted to exhibit increased intestinal carriage duration, resulting in a higher likelihood of onward human-to-human transmission. Yet, bacterial determinants of increased carriage duration are unknown. Previous studies analysed small traveler cohorts, with short follow-up times, or did not employ high-resolution molecular typing, and were thus unable to identify bacterial traits associated with long-term carriage.

**Methods:** In a prospective cohort study of 2001 international travelers, we analysed 160 faecal ESBL-Ec isolates from all 38 travelers who acquired ESBL-Ec during travel and subsequently carried ESBL-Ec for at least 12 months after return, by whole-genome sequencing. For 17 travelers, we confirmed the persistence of ESBL-Ec strains through single nucleotide variant typing. To identify determinants of increased carriage duration, we compared the 17 long-term carriers (≥12 months carriage) with 33 age-, sex- and destination-matched short-term carriers (<1 month carriage). Long-read sequencing was employed to investigate ESBL plasmid persistence.

**Results:** We show that in healthy travelers with very low antibiotic usage, extraintestinal pathogenic lineages of *E. coli* (ExPEC) are significantly more likely to persist than other *E. coli* lineages. The long-term carriage of *E. coli* from ExPEC lineages is mainly driven by sequence type 131 and phylogroup D *E. coli*.

**Conclusions:** Although ExPEC frequently cause extra-intestinal infections such as bloodstream infections, our results imply that ExPEC are also efficient intestinal colonizers, which potentially contributes to their onward transmission.

## Introduction

International travel contributes significantly to the spread of extended-spectrum β-lactamase (ESBL) gene positive *Escherichia coli* (ESBL-Ec).^1,2^ Genetically diverse ESBL-Ec are frequently acquired during international travel.^3–5^ Whilst travel-acquired ESBL-Ec typically are lost during travel or within the first month after return (Figure 1A),^6^ ESBL-Ec and ESBL genes have been detected for more than 12 months after return in a proportion of travelers.^1,3–5,7^ ESBL genes can persist through at least two different mechanisms.^8^ Bacterial strains that carry ESBL-encoding genes in their chromosome or on a stable plasmid can persist over time as part of the local microbiome (strain persistence, Figure 1B). In addition, ESBL genes can be located on mobile genetic elements (MGEs) which persist in the microbiome by their ability to transfer between different bacterial hosts (MGE persistence, Figure 1C).

**Figure 1.**
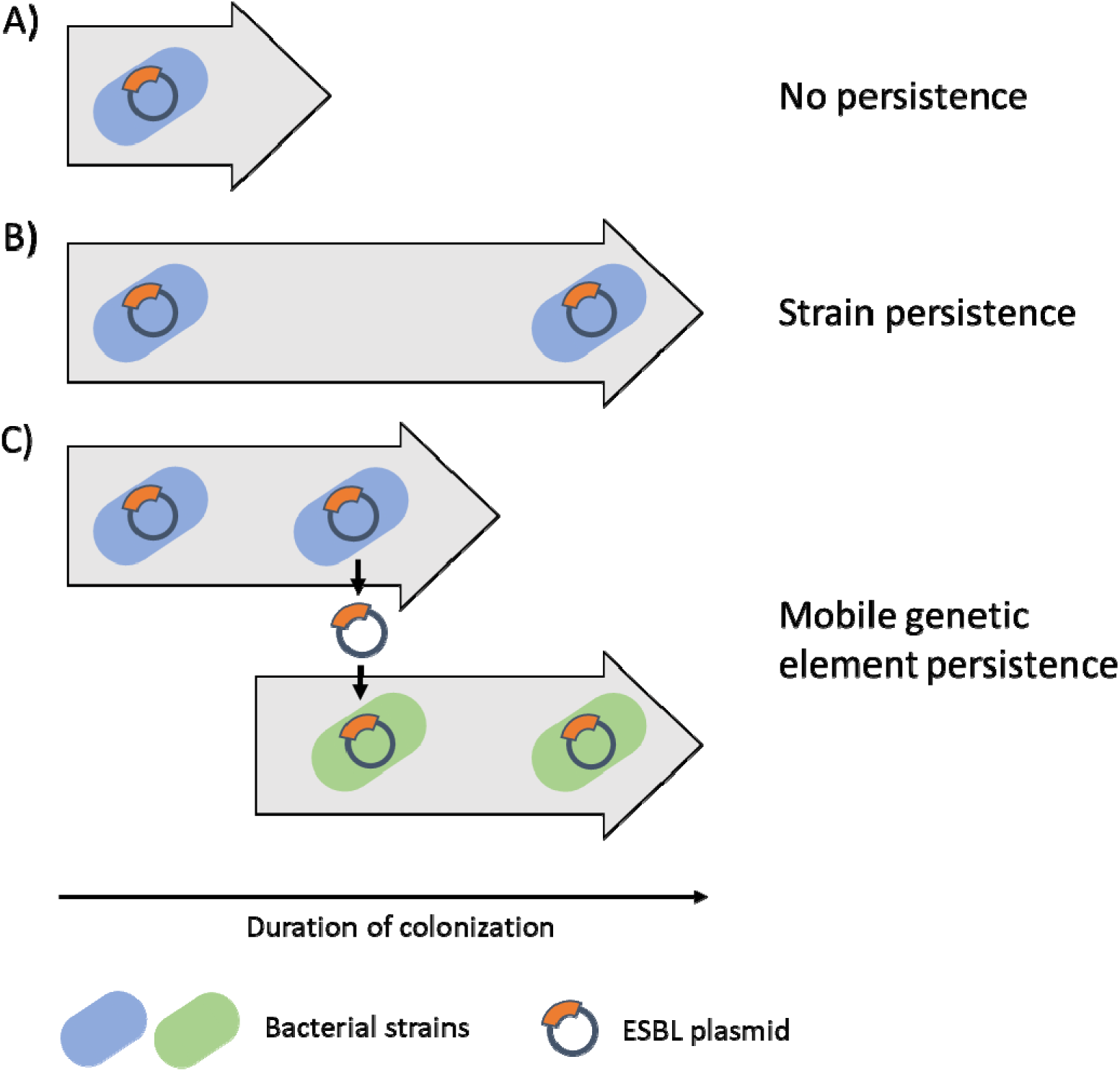
Schematic overview of main persistence mechanism of ESBL genes in the gut of returning travelers. Gray arrows indicate carriage duration over time. A) No persistence, where the colonizing strain is lost during the follow-up period. B) Strain persistence, where a bacterial strain harboring an ESBL gene is continuously present. C) Persistence through mobile genetic elements, where the original strain harboring the ESBL gene has been lost, but the ESBL gene (located on a mobile genetic element) has been passed on to another strain.

ESBL-Ec lineages which are capable of long-term colonization and are adapted to the human intestinal tract are likely to contribute to onward transmission of ESBL-Ec and ESBL genes. However, it is unknown which ESBL-Ec lineages are capable of persistence after return from travel, due to a lack of studies with a sufficiently large sample size and a prospective longitudinal study design. Additionally, high-resolution typing methods such as whole-genome sequencing (WGS) are needed to reliably determine whether the strains have been carried over a long period. One study employed WGS to investigate persistence in 16 travelers who acquired ESBL-Ec abroad and showed that only one traveler carried a travel-acquired ESBL-Ec strain for at least 7 months.^7^ A very recent study analysed data from 11 travellers which were colonised for >3 months.^9^ Due to the low sample sizes in both studies, few ESBL-Ec attributes could be identified that were significantly associated with long-term carriage. Armand-Lefèvre et al. reported an association between phylogroups B2/D/F with prolonged carriage duration (p = 0.02).^9^ Other studies either did not employ WGS or focused on short-term carriage only.

We studied a cohort of 2001 Dutch international travelers (COMBAT cohort).^1^ Six hundred and thirty-three travelers acquired ESBL gene-positive Enterobacterales during travel abroad, of whom 38 travelers (6.0%) were colonized for ≥12 months after acquisition of ESBL gene-positive Enterobacterales, all of which *were E. coli*. Here we report on host and bacterial characteristics associated with ESBL genes persistence and the mechanism through which the ESBL genes persisted. We identified an association between extraintestinal pathogenic *E. coli* and long-term carriage.

## Materials and methods

### Travelers and isolates selection

We included all 38 travelers from the COMBAT cohort (N=2001) who were colonized for ≥12 months with *Escherichia coli* which possessed ESBL genes belonging to a single ESBL gene group at return from travel and at all subsequent time points (1, 3, 6, and 12 months after return), and who tested ESBL-negative before travel (see reference 1 for a detailed description of sampling and microbiological methods).^1^ In short, faecal samples were inoculated in tryptic soy broth containing 50 mg/L vancomycin to select for Enterobacterales. After overnight incubation, the broth was subcultured on chromID ESBL agar plates (bioMérieux). Morphologically different colonies were isolated, with a maximum of five isolates per faecal sample. Eighty-five ESBL-Ec isolates sampled at return from travel (T0) and 75 ESBL-Ec isolates sampled 12 months after return from travel (T12) were included in the current study (Figure 2).

**Figure 2.**
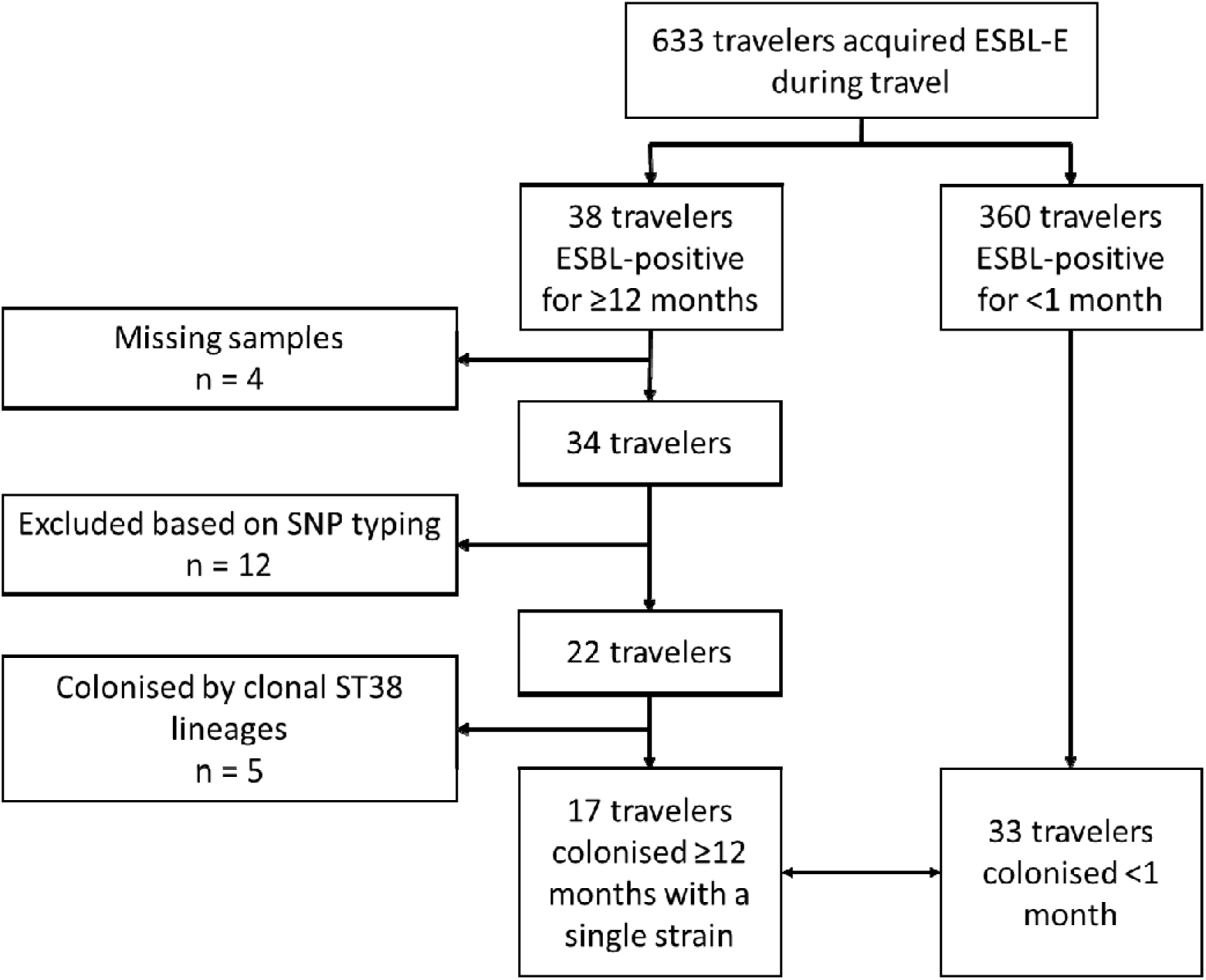
Study flowchart. Strain persistence was defined as isolates from a single traveler, sample twelve months apart with identical ESBL alleles, identical MLST profiles and 25 or fewer genome-wide SNPs difference. Clonal ST38 isolates were excluded. Seventeen long-term carriers (colonized ≥12 months with a single strain) were matched on sex, age and travel destination to thirty-three short-term carriers (colonized <1 month).

### *In silico* typing

DNA extraction and library preparation were performed on all available isolates using the Qiagen Blood and Tissue DNA extraction kit (Cat No./ID: 69506) and Kapa HTP library prep kit (Kit Code KK8234), respectively. Whole-genome sequencing was performed on Illumina HiSeq 2500 by the Amsterdam UMC Core Facility Genomics. Sequencing data were analysed using a Snakemake v5.7.1^10^ pipeline, available at https://github.com/boasvdp/COMBAT (v1.1.0 archived at https://doi.org/10.5281/zenodo.4582689). In short, Illumina sequencing data were trimmed using fastp v0.20.0,^11^ assembled using the Shovill wrapper v1.0.9 (https://github.com/tseemann/shovill) for SPAdes,^12^ and resistance genes were identified using AMRfinderplus v3.2.3.^13^ *E. coli* phylogroups were predicted using EzClermont v0.4.3.^14^ Multi-locus sequence typing (MLST) was performed with the mlst script (https://github.com/tseemann/mlst), using the Achtman scheme for *Escherichia coli*.^15^

### Analysis of strain persistence

We defined strain persistence as the presence of one or more ESBL-Ec isolates in faecal samples from a single traveler obtained at return from travel (T0), as well as 12 months thereafter (T12), with a maximum of 25 SNPs per 5 Mbp difference in the core genome.^16^

To assess strain persistence, we calculated single nucleotide polymorphism differences between isolates to identify which isolates belonged to the same strain. We mapped sequencing reads using Snippy v4.4.5 (https://github.com/tseemann/snippy) on MLST-specific reference genomes selected with ReferenceSeeker v1.6.3^17^ to obtain core genome alignments. Reference genomes used for all MLST sequence types, together with core genome alignment lengths, can be found in table S1. For each MLST, IQtree v1.6.12^18^ was used to infer a phylogeny from the core genome alignment under the model advised by Modelfinder.^19^ Recombination events in the core genome alignment were identified using ClonalFrameML v1.12^20^ and masked using maskrc-svg v0.5 (https://github.com/kwongj/maskrc-svg). SNP differences were counted using snp-dists v0.7.0 (https://github.com/tseemann/snp-dists) and alignment lengths were calculated using a modified version of snp-dists v0.7.0 (https://github.com/boasvdp/snp-dists). SNP counts were scaled to 5 Mbp, to approximate the number of genome-wide SNPs.^16^

To determine whether ESBL plasmids had persisted independent of bacterial host, we employed long read sequencing (Oxford Nanopore Technologies). We generated Oxford Nanopore Technologies sequencing data according to Van der Putten et al. (2020).^21^ In short, strains were grown overnight at 37 °C in liquid LB. DNA extraction and library preparation were performed using the Qiagen MagAttract HMW DNA Kit (Cat. No. 67563) and ONT native barcoding kit (Cat. No. EXP-NBD114), respectively. The library was subsequently sequenced on an ONT MinION flowcell. Raw read data was filtered using Filtlong v0.2.0 (https://github.com/rrwick/Filtlong) and assembled with corresponding Illumina data using Unicycler v0.4.8.^22^ Quality control was implemented at several steps in the pipeline using FastQC v0.11.8,^23^ Quast v4.6.3,^24^ and MultiQC v1.6.^25^ Plasmid comparison was performed using ANICalculator.^26^

### Comparison with short-term carriers

Seventeen long-term carriers were matched by age (range +/-7 years), sex, and travel destination (United Nations subregions) to thirty-three travelers who were colonized for less than a month using SPSS 26 (Figure 2). Illumina WGS was performed as described before on all ESBL-Ec isolated at return from travel from these short-term carriers (total 41 isolates).

### Plotting and statistical analysis

Data were plotted using ggplot2 v3.1.1,^27^ ggthemes v2.4.0,^28^ and patchwork^29^ in R v3.5.1. Tabular data were analysed using Pandas v0.24.2^30^ in Python v3.6.7. Statistical analysis was performed using Fisher’s exact test as implemented in the Python library SciPy.^31^

## Results

### Data characteristics

Out of 2001 Dutch international travelers, we included 34 travelers with samples available, out of all 38 travelers whose faecal samples were positive for *E. coli* harbouring the same ESBL group gene at return (T0), one month after return from travel (T1) and 12 months after return from travel (T12), but were ESBL-negative before travel (Figure 2).^1^ We included a median of 1 ESBL-Ec isolate per traveler per timepoint (range: 1-5 isolates). SNP typing showed that acquired ESBL-Ec isolates were genetically diverse, with some travelers acquiring up to four distinct strains.

### ESBL-Ec strain persistence

#### SNP analysis

Twenty-two out of 34 travelers harboured persistent strains based on SNP typing (Figure 2). Of the 12 travelers who did not harbour persistent strains, ten carried strains with differing MLST profiles between T0 and T12, and two carried strains from a single ST but with the number of core genome-wide SNP differences exceeding the threshold (25 SNPs for a 5 Mbp genome).^16^

#### Clonal ST38 lineages

Non-persistent, yet highly conserved strains that are widely disseminated within the human population could potentially be misidentified as persistent when using SNV analyses. Hence, we compared SNV distances between isolates obtained from unrelated travelers (i.e., belonging to different households) to identify strains shared across our study population. Conserved strains belonging to two ST38 lineages were identified in five unrelated travelers. Lineage ST38-bla_*CTX-M-27*_ was identified in three unrelated travelers and lineage ST38-bla_*CTX-M-14*_ in two other, unrelated travelers. All five unrelated travelers returned from different countries and continents, suggesting these two lineages have disseminated widely. Whole genomes from lineages ST38-bla_*CTX-M-27*_ and ST38-bla_CTX-M-14_ were also abundantly present in public data and a phylogenetic analysis of their core genomes confirmed their high similarity (Figure S1). Hence, we could not conclusively determine whether these highly clonal strains had persisted in the five travelers concerned, or whether these strains were re-acquired from an unknown source. Therefore, we excluded the travelers harbouring these clonal ST38 strains from further analyses to identify lineages potentially associated with persistence. ST38 strains colonizing four additional travelers were not related to the ST38-bla_*CTX-M-27*_. and ST38-bla_*CTX-M-14*_ lineages and were thus not excluded from further analyses.

We identified two related travelers who harboured highly similar ST69 isolates (9 genome-wide SNPs). The most likely explanation of this observation is that the travellers acquired this strain from the same source or one from the other, and that it has persisted in both travellers since.

#### Plasmid analysis

For six travelers, we detected identical ESBL genes from isolates sampled at return from travel and twelve months thereafter, but we could exclude ESBL gene persistence in persistent isolates based on SNP typing (Table S1). To determine whether the plasmid carrying these ESBL genes had persisted independent of bacterial host (Figure 1B), we employed long read sequencing. For one traveler, we detected a pair of ESBL-Ec isolates belonging to different MLST types in which an almost identical plasmid (99.8% nucleotide identity) was identified at T0 and T12, suggesting ESBL gene persistence by plasmid transfer between different *E. coli* hosts.

In summary, we identified persistent ESBL-Ec strains in 17 out of 34 travelers with prolonged ESBL-Ec carriage and persistent ESBL-plasmid in one traveler (Table S1). Although we included multiple ESBL-Ec isolates for some travelers, we did not observe multiple persistent strains within any single traveler.

### Comparison of ESBL-Ec from long-term and short-term carriers

For each long-term carrier (≥12 months carriage) harbouring a persistent strain, two short-term carriers (<1 month carriage) were matched by age, sex and travel destination. For one long-term carrier, only a single matching short-term carrier could be identified, resulting in a comparison of 17 isolates from 17 long-term carriers and 42 isolates from 33 matched short-term carriers, which were sequenced.

Antibiotic usage was low before, during, and after travel and similar between long-term and short-term carriers (Table 1). None of the travelers were admitted to the hospital during or after travel in either group. One single traveler returned to the same country as visited during index travel, within 12 months after return from index travel.

**Table 1.** Characteristics of long-term and short-term carriers. Matching was performed on sex, age and travel destination (United Nations subregions). These characteristics are depicted in italic. Data are presented as number (%) for all characteristics except age and travel duration, which is presented as median (IQR).

|  |  | **Long-term carriers**<br>**(n = 17)** | **Short-term carriers**<br>**(n = 33)** |
| --- | --- | --- | --- |
| *Male* |  | 4 (23.5%) | 8 (24.2%) |
| *Age (years)* |  | 50.2 (41.7-59.4) | 51.4 (36.5-58.1) |
| *Continent*<br>*visited during*<br>*index travel* | *Asia* | 16 (94.1%) | 31 (93.9%) |
|  | *North and South America* | 1 (5.9%) | 2 (6.1%) |
| Travel duration (days) |  | 20 (17-28) | 19 (14-21) |
| Admitted to hospital during index travel |  | 0 (0%) | 0 (0%) |
| Travel to the same country as index travel, within 12 months after return from index travel |  | 1 (5.9%) | 0 (0%) |
| Antibiotic<br>usage | Within 3 months before index travel | 0 (0%) | 0 (0%) |
|  | During index travel | 3 (17.6%) | 2 (6.1%) |
|  | Within 1 month after return from index travel | 0 (0%) | 0 (0%) |
|  | Within 1 to 3 months after return from index travel | 1 (5.9%) | 2 (6.7%) |
|  | Within 3 to 6 months after return from index travel | 0 (0%) | 2 (7.1%) |
|  | Within 6 to 12 months after return from index travel | 3 (17.6%) | 2 (7.4%) |
| Traveler's diarrhoea |  | 6 (35.3%) | 9 (27.3%) |

Following the recent definition of common extraintestinal pathogenic E. coli (ExPEC) lineages according to Manges *et al*. (2019),^32^ persistent strains belonged significantly more often to ExPEC lineages than non-persistent strains. In 15 out of 17 long-term carriers, ExPEC were identified as the persistent strain while in only 7 out of 33 short-term carriers ExPEC were detected (odds ratio 27.86, 95% confidence interval: 5.11-151.74). This difference appears to be driven mostly by ST131 and phylogroup D strains.

## Discussion

We demonstrated long-term carriage of travel-acquired ESBL-positive *Escherichia coli* in 17 travelers out of a cohort of 2001 travelers, which was driven by persistence of ESBL-Ec belonging to ExPEC lineages. In a single traveler, the persistence of ESBL-Ec seemed to be due to an ESBL plasmid which shifted between bacterial hosts after colonising the traveler’s gut. Our study strengthens the finding of Armand-Lefèvre et al.^9^ that ESBL-Ec lineage is associated with persistent carriage after acquisition during travel. Interestingly, we come to similar conclusions although we have focused on long-term carriage (≥12 months) as compared to the previous study (>3 month carriage). Earlier work could not make strong assertions about the persistence of strains, either due to the limited number of included travelers, because the typing methods employed were insufficiently discriminating between isolates, because of a limited duration of follow-up, or because the study population was not representative of community exposure.^3–5,7^ In a recent study in Laos, a group of travelers attending a course at local hospitals acquired a very high diversity of ESBL-Ec E. coli immediately upon arrival and it was shown that this acquisition of resistant E. coli during travel is highly dynamic.^6^ However, the short follow-up in this study did not allow analysis of long-term outcomes of the acquisition of ESBL-Ec.

Previous studies have shown that a limited number of *E. coli* lineages contribute to a large fraction of *E. coli*-mediated extraintestinal disease, including urinary tract infections and bloodstream infections.^32^ These extraintestinal pathogenic *E. coli* are commonly referred to as ExPEC. While ExPEC display pathogenic potential, epidemiological studies often find ExPEC colonizing the human gut in the absence of symptoms.^33,34^ In fact, it has been proposed that ExPEC have evolved towards particularly efficient intestinal colonization and their extraintestinal virulence is an evolutionary “byproduct”.^35^ The long-term persistence of ExPEC lineages, acquired after relatively short travel duration as observed in our study (median 20 days, interquartile range: 17 – 29 days) suggests that ExPEC lineages have spread globally successfully due to their adaptation to the human intestinal tract. It should however be noted that we restricted our analysis to ESBL-producing *E. coli* and duration of persistence of susceptible ExPEC in this cohort is unknown.

Whole-genome sequencing allowed us to detect two clonal ST38 lineages which were shared between unrelated travelers. By additional analysis of publicly available WGS data, these ST38 lineages were shown to have spread globally. The extremely high degree of similarity within these ST38 lineages interfered with reliable strain identification based on core genome DNA sequence analysis. Future studies should explore the application of SNP typing using the pangenome rather than only the core genome, for example through software like Pandora^36^. These novel methods however currently display a higher error rate than the core genome analysis approach used in the present study^36^. Awareness of the circulation of ST38 or other rapidly expanding lineages that are indistinguishable by their core genome is important, for example for management of suspected outbreaks of ST38 in hospital settings.

Our current study focused solely on ESBL-producing *E. coli* and found an association between ExPEC carriage and increased carriage duration. Our approach could also be applied to non-ESBL-producing ExPEC, to determine whether acquisition of ExPEC in general, independent of their antibiotic susceptibility profile, is more likely to results in increased carriage duration after travel. The frequency of sampling can be considered a limitation of this study. However, when comparing between two timepoints, twelve months apart, we could still identify very closely related isolates in samples from the same traveler, indicating strain persistence. Additionally, we only had isolates available for household members of four long-term carriers. Including more household members in future studies might allow us to estimate the likelihood of onward transmission following long-term carriage.

Applying genomic epidemiology to a large traveler cohort, we have shown that ESBL-positive *E. coli* acquired during travel are able to persist for more than a year. The strains that showed the longest carriage duration belonged predominantly to pathogenic ExPEC lineages. Our data imply that long-term carriage of resistant *E. coli* is governed by bacterial characteristics which are associated with lineage. This finding possibly allows a more precise risk assessment for international travelers returning with travel-acquired resistant *E. coli*.

## Acknowledgements

The authors would like to thank Arie van der Ende for the helpful discussions. We thank SURFsara (https://www.surfsara.nl) for the support in using the Lisa Compute Cluster.

## Data and code availability

All Illumina and Oxford Nanopore Technologies sequencing data used in this study are currently available free of restrictions at NCBI under project accession number PRJEB40103. Metadata linking isolates to travelers, required to reproduce our analyses, are currently available free of restrictions at the GitHub repository of this project (https://www.github.com/boasvdp/COMBAT, v1.1.0 archived through Zenodo at https://doi.org/10.5281/zenodo.4582689).

All code is available free of restrictions under the MIT license at https://www.github.com/boasvdp/COMBAT (v1.1.0 archived through Zenodo at https://doi.org/10.5281/zenodo.4582689).

## Ethical statement

The COMBAT study was approved by the Medical Research Ethics Committee, Maastricht University Medical Centre (METC 12-4-093). All participants provided written informed consent.

## Appendices

Figure S1: Phylogeny of clonal complex 38 genomes with continent of origin.

Table S1: All SNP comparisons between isolates with relevant metadata (169 rows). Supplemental information: Members of the COMBAT consortium.

---

**Figure S1.**Core genome phylogeny of 1805 clonal complex 38 *E. coli* strains. Outer ring indicates continent on which the strain was isolated. The two clonal ST38 lineages present in the COMBAT collection are marked in red and blue. Tree and metadata available through iTOL: https://itol.embl.de/tree/2131278347268071589210657.

