## Supplementary figures and images for "Extraintestinal pathogenic *Escherichia coli* (ExPEC) are associated with prolonged carriage of extended-spectrum β-lactamase-producing *E. coli* acquired during travel"

### Figure S1

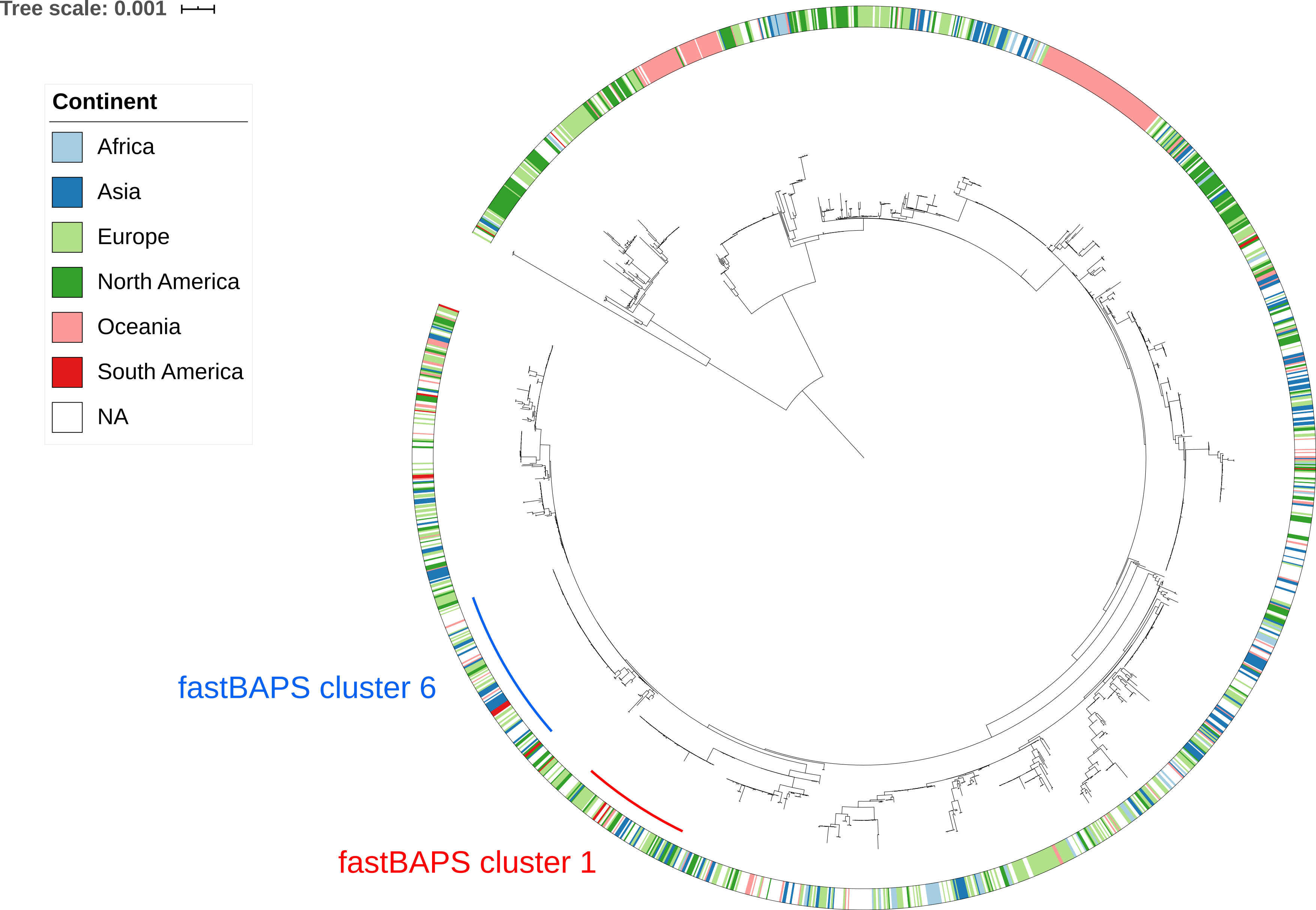
